# Modeling the Sequence Dependence of Differential Antibody Binding in the Immune Response to Infectious Disease

**DOI:** 10.1101/2022.11.30.518471

**Authors:** Robayet Chowdhury, Alexander T. Taguchi, Laimonas Kelbauskas, Philip Stafford, Chris Diehnelt, Zhan-Gong Zhao, Phillip C. Williamson, Valerie Green, Neal W. Woodbury

## Abstract

Past studies have shown that incubation of human serum samples on high density peptide arrays followed by measurement of total antibody bound to each peptide sequence allows detection and discrimination of humoral immune responses to a wide variety of infectious disease agents. This is true even though these arrays consist of peptides with near-random amino acid sequences that were not designed to mimic biological antigens. Previously, this immune profiling approach or “immunosignature” has been implemented using a purely statistical evaluation of pattern binding, with no regard for information contained in the amino acid sequences themselves. Here, a neural network is trained on immunoglobulin G binding to 122,926 amino acid sequences selected quasi-randomly to represent a sparse sample of the entire combinatorial binding space in a peptide array using human serum samples from uninfected controls and 5 different infectious disease cohorts infected by either dengue virus, West Nile virus, hepatitis C virus, hepatitis B virus or *Trypanosoma cruzi*. This results in a sequence-binding relationship for each sample that contains the differential disease information. Processing array data using the neural network effectively aggregates the sequence-binding information, removing sequence-independent noise and improving the accuracy of array-based classification of disease compared to the raw binding data. Because the neural network model is trained on all samples simultaneously, the information common to all samples resides in the hidden layers of the model and the differential information between samples resides in the output layer of the model, one column of a few hundred values per sample. These column vectors themselves can be used to represent each sample for classification or unsupervised clustering applications such as human disease surveillance.

**Author Summary:** Previous work from Stephen Johnston’s lab has shown that it is possible to use high density arrays of near-random peptide sequences as a general, disease agnostic approach to diagnosis by analyzing the pattern of antibody binding in serum to the array. The current approach replaces the purely statistical pattern recognition approach with a machine learning-based approach that substantially enhances the diagnostic power of these peptide array-based antibody profiles by incorporating the sequence information from each peptide with the measured antibody binding, in this case with regard to infectious diseases. This makes the array analysis much more robust to noise and provides a means of condensing the disease differentiating information from the array into a compact form that can be readily used for disease classification or population health monitoring.

## Introduction

Over the past decade, the Johnston lab and others have developed the use of high density quasi-random peptide arrays as a tool for generating antibody binding profilies(4-19). A key feature of these arrays is that the peptide sequences are chosen to cover sequence space as evenly as possible, rather than focusing on biological sequences or known epitopes. Due to the random nature of the peptide sequences, this “immunosignature” approach captures mostly low to moderate affinity interactions of antibodies with the array peptides and has been shown to enable robust detection or identification of immune responses associated with numerous infectious and chronic diseases(8-10, 12-14, 17). This method involves applying a small amount of diluted serum to a dense array of peptides with nearly random sequences of amino acids, typically with >100,000 distinct peptide sequences of about 10 amino acids in length(7). In most of the studies done, only 16 of the 20 natural amino acids were used to synthesize the peptides. The level of antibody binding to the peptides on the array is then detected quantitatively using a fluorescently labeled secondary antibody and imaged by an array scanner. Based on a statistical comparison of binding patterns between case and reference samples, classifier models can be built to distinguish one disease response from another(5).

The cognate epitopes of the antibodies involved in an immune response are highly unlikely to appear within a random set of ∼10^5^ sequences on a peptide array. For a linear epitope of ∼10 amino acids in length, there are ∼10^13^ possible amino acid combinations, yet somehow the interaction of serum antibodies with only ∼10^5^ sequences captures sufficient information to both detect and identify disease state with high accuracy(6-10, 12-14, 17, 20). If sufficient information can be obtained from a random sparse sampling of antibody binding to 1 out of every 10^8^ possible sequences (∼10^13^/∼10^5^), then the antibodies associated with an immune response must recognize millions to billions of different sequences to some extent in a manner that is disease specific. The fundamental question of the current study is whether this amino acid sequence-dependent antibody binding can be modeled. If so, such a relationship could potentially be used to more effectively aggregate information from the array or to design new panels of sequences that more effectively differentiate diseases.

Recently, our group modeled the sequence-binding relationships of nine different, well-characterized, isolated proteins to the peptide arrays described above(21). Binding patterns of each protein were recorded, and a simple feed-forward, back propagation neural network model was used to relate the amino acid sequences on the array to the binding values. Remarkably, it was possible to train the network with 90% of the sequence/binding value pairs and predict the binding of the remaining sequences with accuracy equivalent to the noise of the antibody binding measurements (the Pearson correlation coefficients (R) between the observed and predicted binding values were equivalent to that between measured binding values of multiple technical replicates, and in some cases as high as R=0.99). In fact, accurate binding predictions (R > 0.9) for some protein targets could be achieved by training on as few as hundreds of randomly chosen sequence/binding value pairs from the array. In addition, the binding predictions were specific; the model captured not only the bulk binding of individual proteins but also the differential binding between proteins. Finally, a neural network trained on weakly binding sequences effectively predicted the binding values of sequences on the array 1-2 orders of magnitude greater. At least in the context of the combinatorial space of possible sequences in this model array-based system (∼10 residue peptides using 16 different amino acids with the C-terminus bound to the surface of a silica substrate), training on one set of thousands of randomly selected sequences resulted in statistically accurate prediction of the binding to any other randomly selected set of sequences.

Binding to antibodies, in this case IgG in human sera, represents a much more complex system than binding to isolated proteins, and one might expect substantially more complex sequence-binding relationships. Other groups have previously developed such relationships for immune responses using various starting datasets. A number of groups have looked at overlapping peptides presented on microarrays or in phage display libraries generated by tiling antigens or entire proteomes(22-27). Panning of phage or bacterial peptide display libraries coupled with next generation sequencing have provided broader binding profiles(28, 29). The advantage of tiling and panning approaches is that one is starting with known or suspected binding sequences, and thus the dataset is naturally rich in strong binding information. In one particularly effective study in this regard, a method referred to as Protein-based Immunome Wide Association Study was used to explore sequence binding relationships in 31 systemic lupus erythematosus samples(30). Here a large bacterial display library (10^10^ 12-mer sequences) was reduced to ∼10^6^ sequences found to bind to serum antibodies from the samples and the enrichment of specific 5-mer and 6-mer sequences within the resulting library was determined. These enriched sequences were then used to identify autoantibodies in the human proteome, and the authors were successful at identifying several known autoantigens for the disease within their top candidates. The same group has used similar methods to perform epitope mapping of antibodies to SARS-CoV-2(31).

Machine learning algorithms have also been used to develop sequence-based models predicting binding of proteins to peptides, antibodies, and DNA(32-42). For example, machine learning models have been used to model anti-microbial peptides, infectious viral variants that escape protection, potential epitopes on target antigens, high antibody binding regions on target proteins, and optimization of target DNA sequences for transcription factors. To do this, two approaches have primarily been used: 1) introducing single or multiple point mutations on a target site with known function to identify desired leads, and 2) use of proteomes of interest or known antigenic proteins to predict epitopes. For example, epitope prediction tools such as BepiPred-2.0 are generally developed using known antigens derived from crystal structures of antibody-antigen complexes(43). With regard to modeling of serum binding to random sequences, Greiff *et al*, applied multivariate regression to serum antibody binding to a library of 255 random peptides(44). In that study, serum antibody binding from naïve mice was well modeled by relating peptide composition to binding intensity, though binding of serum antibodies from previously infected mice proved more challenging.

The current work focuses on the feasibility of developing comprehensive sequence-binding relationships that describe the infectious disease specific binding of total IgG to our model library of 122,926 peptides each between 7 and 12 residues in length and composed of 16 of the 20 natural amino acids. While this library is clearly limited in terms of size (only 10^5^ of the trillions of possible sequences), composition (16 of 20 natural amino acids) and context (C-terminus affixed to a silica surface), it is capable of distinguishing immune responses to different infectious agents, as described previously(6-8, 13). Neural network-based models were used to build quantitative relationships for sequence-antibody binding using sera from cohorts of individuals who are either uninfected (controls) or infected with 5 infectious agents including three closely related members of the family *Flaviviridae* (dengue virus, West Nile virus and hepatitis C virus), a more distantly related member of the family *Hepadnaviridae* (hepatitis B virus) and an extremely complex eukaryotic trypanosome (agent of Chagas disease, *Trypanosoma cruzi*). This allowed a thorough evaluation of the model’s ability to capture the disease-specific information content of the array binding. This study showed that it is possible to create accurate sequence-binding models, which not only maintain the disease specific information, but also effectively capture the binding information on the arrays for applications in noise suppression and disease classification.

## Results

### Study Design and Initial Analysis

The serum samples shown in Table 1 were incubated on identical peptide microarrays as described in Methods and IgG bound to the array peptides was detected via subsequent incubation with a secondary anti-IgG antibody. The peptide sequence ‘QPGGFVDVALSG’ is present on the array as a set of replicate features (n=276). This peptide sequence gives a consistently moderate to strong binding value from sample to sample and is used to assess the intra-array spatial uniformity of antibody binding intensities. Median normalized arrays with an intra-array replicate feature coefficient of variation (CV) ≥ 0.3 for this peptide sequence were set aside as well as arrays that showed significant physical defects or overall differences in binding intensity between different regions of the array (collectively these are referred to as “High CV samples”). In all, 20% of the 679 arrays measured were excluded from the initial part of the analysis but considered in the last section which focuses on using the sequence-binding relationship to remove noise from the arrays. Thus, 542 arrays total were considered “Low CV Samples” in Table 1.

**Table 1:** Sample information.

| Disease cohort | Sample Source<sup>1</sup> | Samples Collected | Low CV Samples<sup>2</sup> | Genome Size(bp) |
| --- | --- | --- | --- | --- |
| Hepatitis C Virus (HCV) | CTS | 100 | 78 | 11,000 |
| Dengue Virus, Serotype 4 (Dengue) | CTS and SeraCare | 65 | 57 | 9600 |
| West Nile Virus (WNV) | CTS | 100 | 74 | 11,000 |
| Hepatitis B Virus (HBV) | CTS | 100 | 86 | 3200 |
| *T. cruzi* | CTS | 96 | 70 | 105M |
| Uninfected (ND) | CTS and ASU | 218 | 177<sup>3</sup> | -- |

<sup>1</sup>CTS is Creative Testing Solutions (Tempe, AZ); ASU is Arizona State University; SeraCare address is Milford, MA
<sup>2</sup>Arrays passing the data quality metrics used in the initial neural network analysis. The remaining high CV samples were used as a test set for certain classification studies.
<sup>3</sup>100 randomly selected uninfected samples were used for the bulk of the neural network analysis to remain reasonably balanced with other cohorts.

#### Comparison of average binding profiles of peptides to serum IgG

Figure 1 shows the cohort average serum IgG binding intensity distributions of the 122,926 unique peptide sequences. The samples were all median normalized prior to averaging each peptide binding value within the cohort. The log_10_ of the average binding is displayed on the x-axis as the log distributions are much closer to a normal distribution than are the linear binding values. Sera from individuals infected with HCV, dengue virus or WNV have sharper distributions (smaller full width at half maximum) than the other samples, while sera from individuals infected with HBV show a distribution width similar to those from uninfected donors. Sera from individuals with Chagas disease have a broader binding distribution than the others, with a long tail on the high binding side. Overall, the width of the distribution increases with increasing proteome size. Interestingly, for the viruses with small proteome some of the higher binding antibodies are lost compared to uninfected samples. However, it is important to remember that the array peptides have no relationship to the viral proteomes or indeed any biological proteome, except by chance. Thus, what is lost in the small virus samples compared to strong binding in uninfected samples, may well be gained in more specific binding not immediately apparent.

**Fig. 1.**
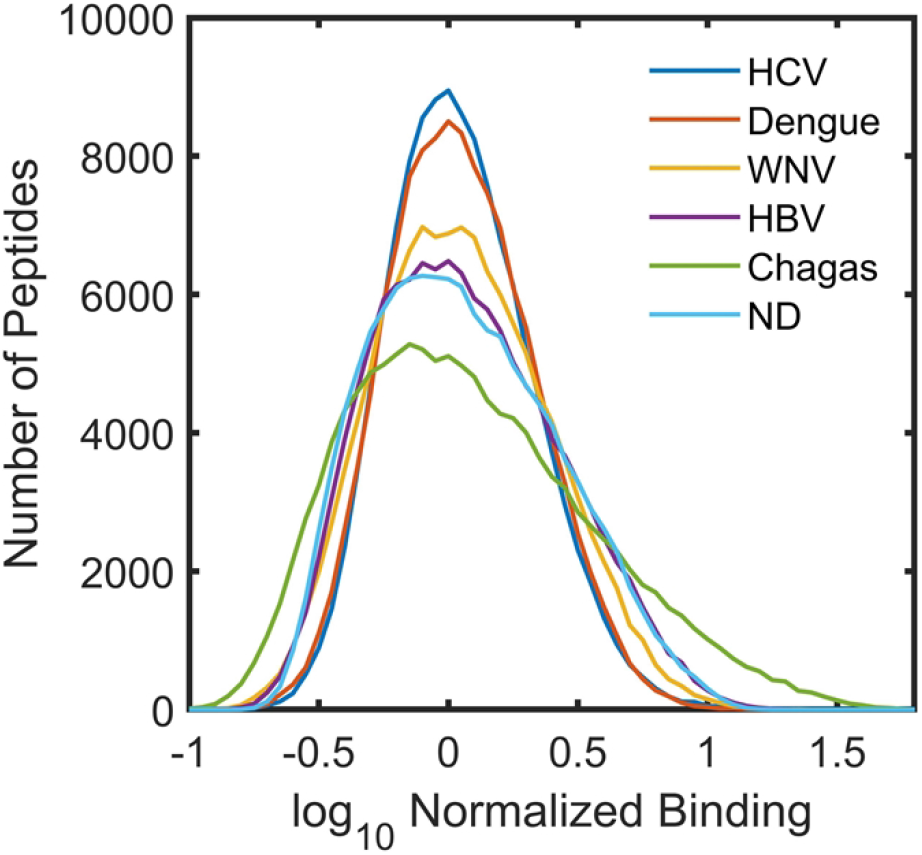
Average Binding Distributions of the Cohorts. Average binding intensity distributions of serum IgG binding to array peptides for the 6 different sample cohorts. For each cohort the log_10_ of the average binding for each peptide sequence was used to create the distribution.

### Neural Network Analysis

The fundamental question of this study is whether it is possible to accurately predict the sequence dependence of the antibody binding associated with an immune response to a given pathogen, both in terms of accurately representing the IgG binding to each peptide sequence in individual serum samples and in terms of the ability of the neural network to capture sequence dependent differences in IgG binding between samples and cohorts. Towards this end, the low CV samples (Table 1) were analyzed using feed forward, back propagating neural network models(21) in two different ways. In one approach, each sample was analyzed separately such that a neural network was trained on every serum sample independently. In the second approach, all samples were fit together such as that a single neural network was trained to simultaneously predict the binding for all samples for any given sequence. In both cases, the optimized network involved an input layer with an encoder matrix (see Methods), two hidden layers with 350 nodes each and an output layer whose width corresponded to the number of target samples (1 for individual fits and 465 when all samples were fit simultaneously). The loss function used was the sum of least squares error based on a comparison of the predicted and measured values for the peptides in the sample.

#### The neural network uses the sequence information to rapidly converge on a solution

Fig. 2A shows the rate at which the loss function drops during training using the simultaneous fitting approach in which all samples are analyzed together. When the correct sequence is paired with its corresponding binding value (blue and red lines, Fig. 2A), the value of the loss function drops rapidly and the values for the training set and test set drop in concert; there is almost no overfitting. As a control, the same neural network was used to analyze data in which the order of the peptide sequences was randomized relative to their binding intensities. One would not expect any relationship between sequence and binding under these circumstances. In this case, the loss function value for both the training and test initially rise slightly followed by a slow drop for the training set of peptides over the entire training period and a slow rise for the test set (yellow trace: test, purple trace: train) indicating overfitting of the training set. This implies that the neural network is capable of rapidly converging on a true relationship between the sequences and their binding values in the context of the array peptide library.

**Fig. 2.**
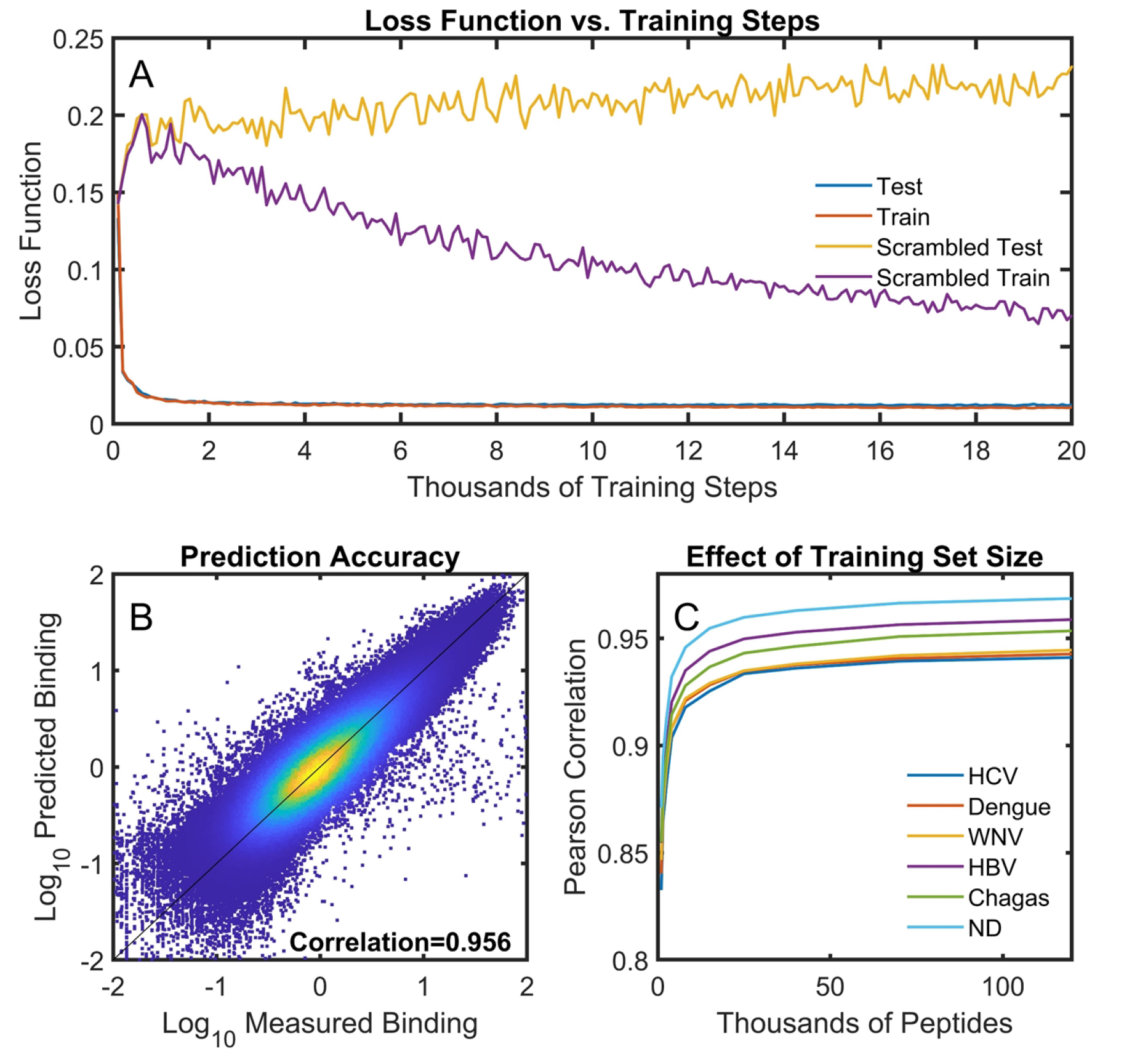
The neural network model accurately represents the sequence-binding relationship. A neural network (2 hidden layers with 350 nodes) was trained on 95% of the sequence/binding data from the 465 low CV samples in Table 1 simultaneously (note only 100 of the uninfected samples were used to balance with the size of other cohorts). The remaining 5% of the sequence/binding values (6,146 per sample x 465 samples = ∼2.9 million binding values) were held out as the test set. (A) The loss function progression during neural network training. Blue and red traces (overlapping): a neural network trained with properly matched sequences and associated binding values. Purple and yellow traces: training after scrambling the order of the sequences relative to their measured binding values. (B) The scatter plot (dscatter(2)) shows the values predicted by neural network (y-axis) vs. the corresponding measured values from the array (x-axis) for the test set only. (C) The average predicted vs. measured correlation coefficient for cohort samples as a function of the number of peptide sequences used to train the network.

#### The neural network results in a comprehensive binding model applicable across the model sequence space used

Fig. 2B shows a scatter plot comparing the predicted and measured values from a neural network model fitting all samples simultaneously. In this case, the model was trained on 95% of the peptide sequence-binding pairs, randomly selected, with the remaining 5% or 6,146 peptide sequences excluded from training and used for model testing (that is 6,146 binding values for each of the 465 low CV samples used = ∼2.9 million binding values in the test set). Only the test set values are displayed in Fig. 2B. Since the sequences used on the array are nearly random, these sequences should be statistically equivalent to any randomly selected set of sequences from the combinatorial space of possible sequences sampled by the array (peptides of about 10 residues utilizing any of 16 amino acids corresponds to about 10^12^ sequences). The Pearson correlation coefficient (R) between the measured and predicted values for the test sequences shown is 0.956. Repeating the training 100 times with randomly selected train and test sets gives an average correlation of 0.956 with a standard error of the mean of 0.002. The correlation coefficient between measured and predicted binding for the 95% of the sequences used to train the neural network was 0.963 +/-0.002. This implies that there is almost no overfitting associated with the model (the quality of fit between the test and train data is similar), a conclusion also apparent in the loss function data of Fig. 2A. Fig. S1 shows the correlation coefficient between measured and predicted binding for each individual sample in the test dataset (using a simultaneous fit of all samples). While some cohorts and some samples were better represented than others, for the vast majority of the samples, the correlation coefficients are greater than 0.9.

#### 10^3^ to 10^4^ peptides are sufficient to provide a reasonable description of the entire combinatorial peptide sequence space

Neural network models were trained with different numbers of randomly selected peptides, and binding was predicted for the remaining portion of the peptides. Fig. 2C explores the dependence of the overall correlation coefficient between measured and predicted binding values for the test set of each of the sample cohorts as a function of the number of peptides used in the training. When at least 10,000 peptide sequences are used to train the neural network, the correlation coefficient is >0.9 for all cohorts, and the correlation is >0.85 when the model is trained using only 2,000 peptides. This implies that even a very sparse sampling of this sequence space provides a reasonably accurate model of the sequence-binding relationship. The correlation coefficients do continue to increase slowly as a function of training set size. Thus, even though a relatively small set of peptides gives a reasonable overall picture, the predictive power of the relationship continues to improve with more data, and if even more peptide sequences were available for training than the entire 122,926 peptides on the array, an improved prediction would be expected.

#### There are commonalities in the binding of each sample that make simultaneous modeling of all samples more accurate than individual neural network models

As stated above, it is possible to either build entirely independent neural network models for each of the samples considered or to build models that fit all of the samples simultaneously. Fig. 3 shows a direct comparison of the measured vs. predicted correlation coefficients of each sample using the simultaneous and individual model approaches. In almost every case, the simultaneous model is more accurate, providing a small improvement in correlation coefficient. This implies that the network learns commonalities between IgG binding from serum across all samples and different cohorts and uses those commonalities to improve the model. In the simultaneous model, these common features are learned by the 2 hidden layers of the neural network and the differences between samples are learned in the output layer (the final weight matrix), with separate columns in that layer giving rise to the binding values for each sample. Simultaneous modeling of all the samples is used for the remainder of the analyses in this work. Simultaneous modeling was also dramatically faster than fitting each sample dataset separately. For comparison, a simultaneous training required about 10 minutes to complete on an 18 CPU core machine while the individual modeling required about 10 hours even after optimizing parallel processing.

**Fig. 3.**
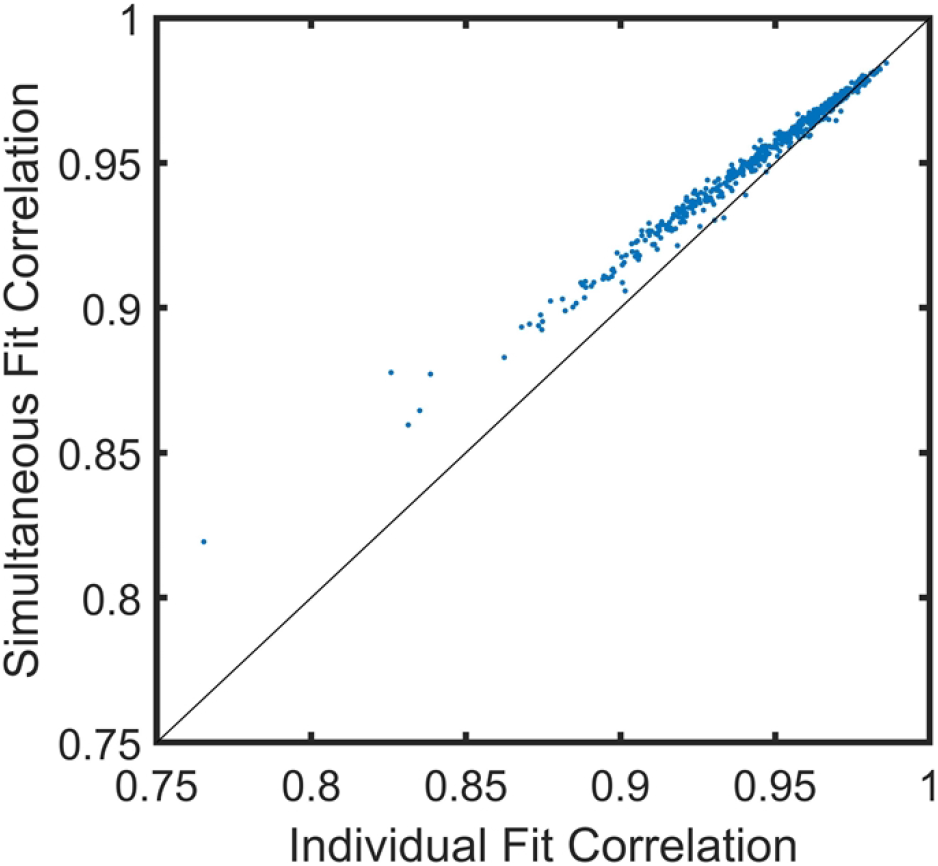
Simultaneous Modeling of All Cohorts is More Accurate Than Individual Fits. A comparison of predicted vs. measured correlation coefficients calculated either by fitting samples simultaneously (as in Fig. 2) or one at a time.

### The Neural Network Learns Distinguishing Characteristics of Cohorts

Fig. 4A is a schematic of three approaches to disease classification and discrimination. The blue line is the standard statistical pathway (immunosignaturing). Here, no sequence information is used in the analysis and the binding values are either fed into a classifier (Fig. 4B) or used to determine the number of significant peptides that distinguish diseases (Fig. 4C), as described below. Alternatively, the neural network can be used to determine a sequence/binding relationship. This relationship can either be used to recalculate predicted binding values for the array peptide sequences, forcing the data to always be consistent with the sequences (red line), or it can be projected onto a completely new set of sequences (an *in silico* array, orange line), and those projected binding values used in classification or determining the number of significant distinguishing peptides between disease pairs.

**Fig. 4.**
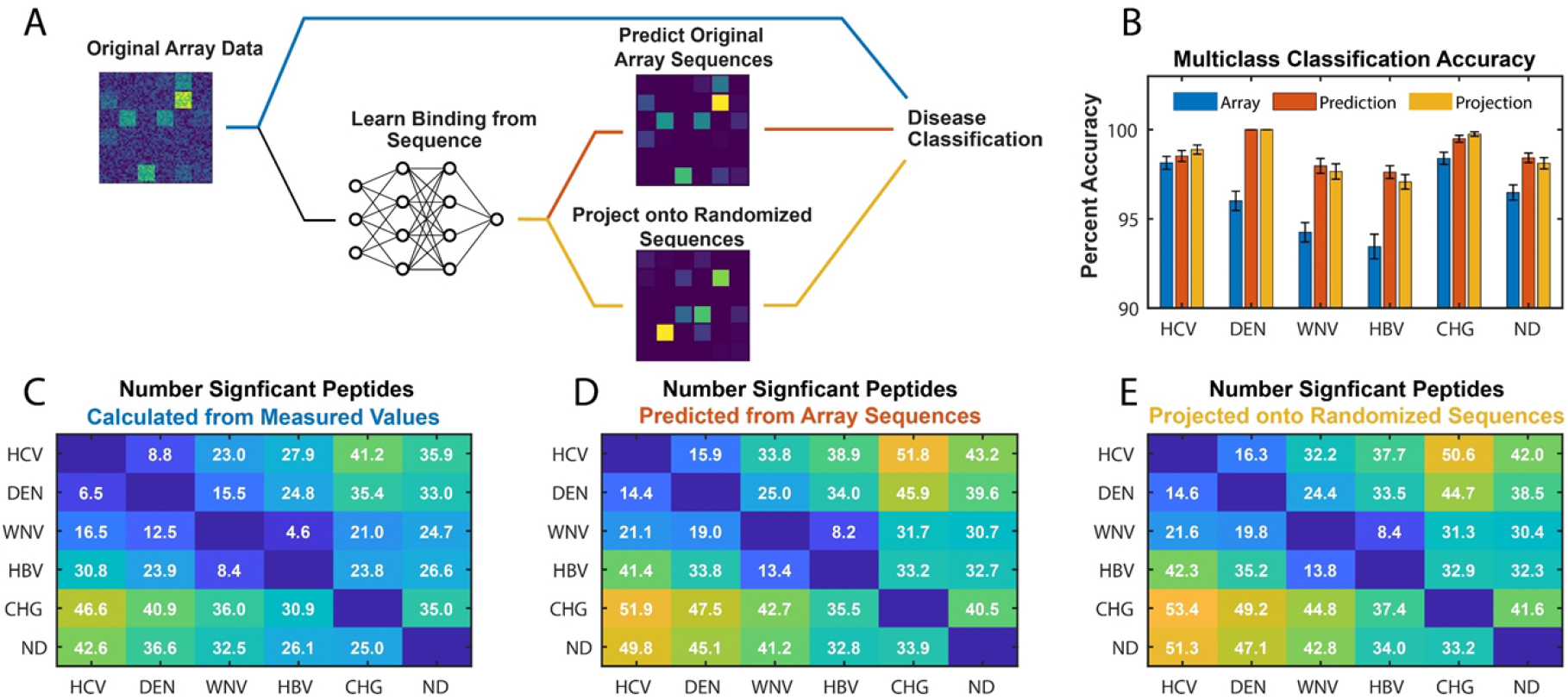
Discriminating between cohorts. (A) The data from the original array was analyzed in three ways: directly (blue line), after training a neural network and predicting the values of the array sequences (red line), and after projecting the trained neural network on a complete new set of sequences (orange line). Disease discrimination was then performed for each approach using multi-class classification or by statistically determining the number of significant peptides distinguishing each cohort comparison. (B) Multi-class classification based on a neural network (see text). Classification was performed 100 times for each dataset leaving out 20% of the samples (randomly chosen) each time. Blue: original measured array data. Red: neural network model prediction of binding values for array peptide sequences. Orange: neural network projected onto a randomized set of sequences of the same overall size, composition and length distribution as the array sequences. (C) Each array element is the number of array peptides with measured binding values that are significantly higher in the sample cohort on the Y-axis compared to the sample cohort on the X-axis. Significance is defined as a p-value less than 1/N in a T-test with 95% confidence (N = 122,926 total peptides, thus significant peptides have a p-value < 0.05/N = 4.1×10-7). (D) As in (A) except that the neural network predicted binding values of the array peptides were used instead of the measured. The mean of 10 different neural network model training runs is shown; error in the mean is ≤0.3. (E) The same as in (D) except predicted values for an *in silico* generated array of random peptide sequences with the same average composition and length as the peptides in the array were used. The mean of 10 different sequence sets and neural network runs is shown; error of the mean is ≤0.4.

#### Values predicted by the neural network result in better ability to distinguish cohorts

In Fig. 4C-E, the number of peptide binding values that are significantly greater in one cohort (on the Y-axis) compared to another (on the X-axis) are shown in each grid. Significance was determined by calculating p-values for each peptide in each comparison using a T-test between cohorts adjusted for multiple hypothesis comparisons using the Bonferroni correction.

Significant peptides are those in which the p-value is less than 1/N (N=122,926) with >95% confidence. Fig. 4C shows comparisons between cohorts using the measured data from the arrays. As one might expect, the sera from donors infected with the Flaviviridae viruses are most similar to one another in terms of numbers of distinguishing peptides. In general, they are more strongly distinguished from HBV (except for WNV) and very strongly distinguished from Chagas donors. If one follows, for example, the top row of Fig. 4C for HCV, moving to the right one sees that the numbers increase as more and more genetically dissimilar comparisons are made. West Nile virus is an exception in this regard. While it is more similar to Dengue virus than it is to Chagas, it is most similar, in terms of numbers of distinguishing peptides, to HBV (Fig. 4C).

Figure 4D is the same as Fig. 4C except that in this case, the predicted values from the neural network model are used for the array sequences instead of the measured values. Because the network requires that a common relationship between sequence and binding be maintained for all sequences, it increases the signal to noise ratio in the system such that significantly more distinguishing peptides are identified in every comparison. The neural network was run 10 times and the results were averaged.

Figure 4E shows results in the same format as the other two panels but using *in silico* generated sequences and their binding values predicted by the neural network model trained on peptide array binding data. These sequences were produced by taking the amino acids at each residue position in the original sequences and randomizing which peptide they were assigned to (considering the sequences as a matrix with rows representing peptides in the array and columns representing residue positions, order of amino acids in each column was randomized separately and at the end any spaces due to varying peptide lengths were removed). This created an *in silico* array with a completely new set of sequences that had the same number, overall amino acid composition and average length as the sequences on the physical array to ensure a consistent comparison. The binding values for each sample were then predicted for this *in silico* array and those values were used in the cohort comparisons. The number of significant peptides identified using the new sequence set (Fig. 4E) are identical to within error for each comparison with the predictions from the actual array peptide sequences used in the training (Fig. 4D). Note that the result of generating ten different randomized *in silico* arrays was averaged.

Another way to understand how well distinguishing information is captured by the neural network model is to compare classification based on measured values *vs*. predicted values. Fig. 4B shows the result of applying a multiclass classifier, either to the measured binding values, the binding values predicted for the array sequences, or binding values predicted for *in silico* generated sequences. A simple multiclass classifier was built using a neural network with a single hidden layer with 300 nodes (described in the supplementary information). This will be referred to simply as the “multiclass classifier” to avoid confusion with the neural network used to model the sequence-binding relationship. The multiclass classifier cannot effectively use all peptides for each sample. Peptide feature selection was performed using a peptide-by-peptide T-test between the binding values of each cohort vs. all others. Either 20 features (the measured data) or 40 features (the two predicted data sets) were used per cohort, with the number of features chosen to be optimal for the dataset (see Fig. 4 caption). The training target is a one-hot representation of the sample cohort identity, and the network is set up as a regression. 80% of the samples were randomly selected and used to train the multiclass classifier and 20% were used as the test set. Each test sample was then assigned a cohort label based on the largest value in the resulting predicted output vector. The process was repeated 100 times and overall prediction accuracy determined. For every cohort, with the possible exception of HCV, classification was improved relative to direct use of the measured array values (blue bars) when using the predicted values. This was true using either predicted values for the array sequences (red bars) or predicted values resulting from projection of the trained network on the randomized *in silico* array sequences (orange bars).

### Understanding the Noise Reduction Properties of Neural Network Modeling

The results presented above show that by using the sequence/binding information to first train a neural network model and then predicting the binding using that model (on the same or a different set of sequences), it is possible to improve the signal to noise ratio in the data, at least for the purpose of differentiating between disease cohorts. To understand this in more detail, the effects of noise added to the data was explored.

#### Gaussian noise is effectively removed by the model

In Fig. 5, noise was artificially added to each point in the measured dataset by using a random number generator based on a gaussian distribution that was centered at the measured value:

**Fig. 5.**
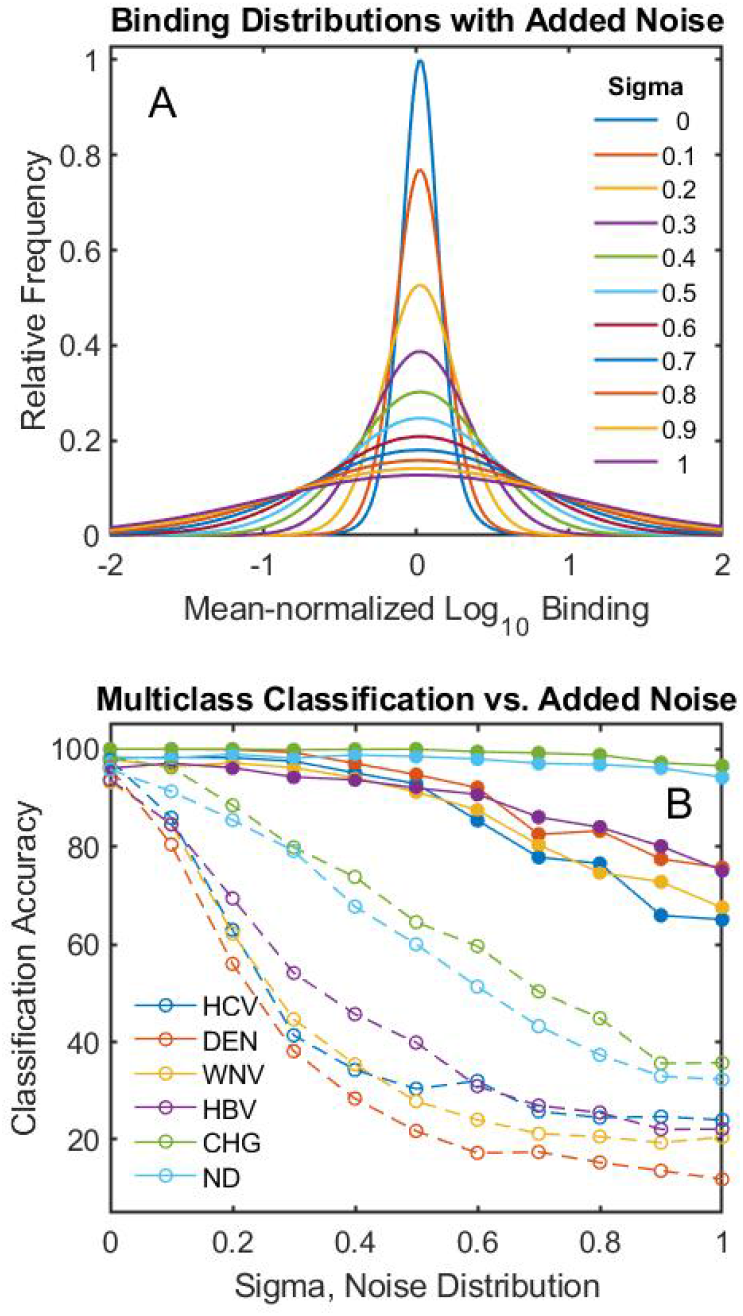
Effect of added noise on multiclass classification. Noise was added to each peptide in the sample using a randomly chosen value from a gaussian distribution centered at the log_10_ of the measured value. The sigma of the distribution was varied between 0 and 1 (the binding, and thus sigma, is on a log scale). (A) The resulting distributions of binding values for each sigma value. Distributions were determined after mean normalizing the binding values for each peptide in a cohort and then including all peptide binding values in the distribution. (B) Results of applying a multi-class classifier (as in Fig. 4B) to the data for measured binding values (dashed lines) and predicted binding values (solid lines) at each value of sigma. Each classification was repeated 100 times (noise at each level was randomly added 10 times and each of these were reclassified 10 times leaving out 20% of the samples as the test set).

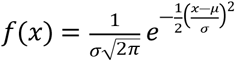

In the above equation, mu (µ) is the log_10_ of the median normalized measured binding value. Sigma (σ) was then varied from 0 to 1 to give different levels of added noise. Note that sigma =1 results in addition of noise on the order of 10-fold greater or less than the linear binding value measured (due to the log_10_ scaling). Fig. 5A shows the resulting distribution of peptide binding values after adding noise. The peptide binding values were mean normalized across all cohorts and then plotted as a distribution, for each cohort (since this is the log_10_ of the mean normalized value, the distributions are centered at 0). As sigma is increased, the width of the resulting distribution after adding noise increases dramatically.

Fig. 5B plots the multi-class classification accuracy of each dataset for each sample cohort as a function of sigma (this uses the same multiclass classifier as Fig. 4). The classification accuracy of the original measured data with increasing amounts of noise added drops rapidly (dashed lines). Since this is a 6-cohort multi-class classifier, random data would give an average accuracy of ∼17%. The measured values with added noise approach that accuracy level at the highest noise. However, by running the data through the neural network and then using predicted values for the same sequences as are on the array, the accuracy changes only slightly for sigma values up to about 0.5 and then drops gradually with increased noise, but always remains well above what would be expected for random noise. Note that a sigma of 0.5 corresponds to causing the linear measured values to randomly vary between about 30% and 300% of their original values.

#### Neural network predictions of array signals improved classification of high CV samples

As described above, 137 samples were not used in the analyses above because they either had high CV values calculated from repeated reference sequences across the array or because there were visual artifacts such as scratches or strong overall intensity gradients across the array. A neural network model was applied to all 679 sample in Table 1 (all 542 low CV + 137 high CV) simultaneously. Note that the model does not include any information about what cohort each sample belongs to, so modeling does not introduce a cohort bias. The overall predicted *vs*. measured scatter plots and correlations are given in Fig. 6A and 6B for the low CV and high CV data, respectively. The number of points displayed was randomly selected to be constant between datasets and make the plots comparable. Prediction of the binding values for the high CV data results in more scatter relative to measured values, due to the issues with those particular arrays.

**Fig. 6.**
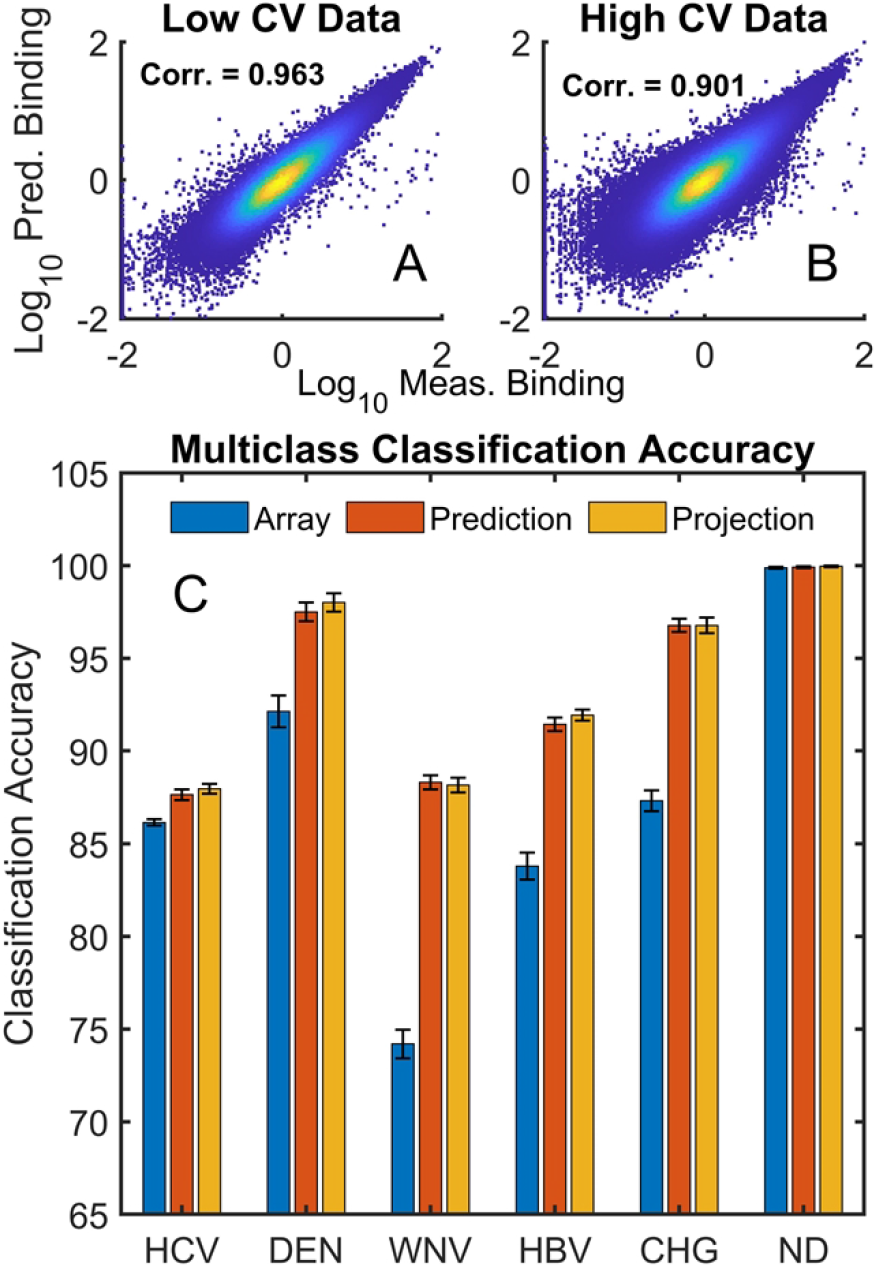
Classification accuracy for high CV samples. (A) Neural network predicted vs. measured values for low CV data and (B) for high CV data. (C) Multiclass classification of the high CV data. Blue, Red and Orange bars represent use of measured, predicted and projected data as in Fig. 4.

In Fig. 6B, the measured and predicted values for the 542 low CV samples were used to train a multiclass classifier which was then used to predict the cohort class of the high CV samples. Three different data sources were used: 1) the measured array data (blue bars), 2) predicted binding values for the array peptide sequences based on the neural network model (red bars) and 3) projected values for *in silico* generated arrays similar to those used in Fig. 4 (orange bars). The classifier used was the same as that in Fig. 4 and the number of features selected was optimized for the data source as described for the analysis of Fig. 4 (20 features per cohort for the measured array data and 40 features per cohort for the two datasets based on the neural network predictions). In each case except for the non-disease samples, the use of predicted values resulted in a significantly better classification outcome.

## Discussion

### A Quantitative Relationship Between Peptide Sequences and Serum IgG Binding

The work described above shows that it is possible to use a relatively simple neural network model to generate a quantitative relationship between amino acid sequence and serum antibody binding over a large amino acid sequence space by training on a very sparse sampling of binding to that sequence space, similar to what was seen previously for isolated proteins binding to the array(21). Indeed, a reasonably accurate prediction can be obtained with only thousands of sequences (Fig. 2C).

The model system used here to explore the relationship between antibody molecular recognition profiles and amino acid sequences has limitations. Only 16 of the 20 natural amino acids were used in this model for technical reasons (see Materials and Methods). The sequences are also bound at one end to an array surface, and the other end has a free amine rather than a peptide bond as would be seen in a protein. In addition, the array peptides are short, linear and largely unstructured. This limits the range of molecular recognition interactions that can be observed, and thus the level of generality of the conclusions, but also suggests that comprehensive and accurate structure/binding relationships for humoral immune responses should be possible to generate given binding data in a broader sequence context. Such relationships would be invaluable for epitope prediction, autoimmune target characterization, vaccine development, effects of therapeutics on immune responses, etc. Even this rather simple model system for sequence space already shows the ability to capture differential binding information between multiple diseases simultaneously, including infectious diseases that involve closely related pathogens (Fig. 4).

The fact that one can develop comprehensive sequence/binding relationships within this model sequence space also explains, at least in part, why the immunosignature technology is promising. Immunosignaturing technology as applied to diagnostics uses the quantitative profile of IgG binding to a chemically diverse set of peptides in an array followed by a statistical analysis and classification of the resulting binding pattern to distinguish between diseases. The approach has been successfully used to discriminate between serum samples from many different diseases (6-10, 12-14, 16, 17) and has been particularly effective with infectious disease(6-8, 18), as exemplified by the robust ability to classify the immune response to the infectious diseases studied here (Fig. 4D). This raises the question, why would antibodies that are generated by the immune system to bind tightly and specifically with pathogens show any specificity of interaction to nearly random peptide sequences on an array? The success of the neural network in comprehensive modeling of the sequence/binding interaction provides an answer. The *information* about disease-specific IgG binding is dispersed broadly across peptide sequence space, even in the interaction with sequences that themselves bind weakly and with low specificity, rather than being focused only on a few epitope sequences. It is not necessary to measure binding to the epitope if you have a selection of sequences that are broadly located in the vicinity of the epitope in sequence space.

Note also that by working with sequence/binding relationships, rather than purely statistical comparisons of binding values associated with specific sequences, one can combine information from arrays that contain different peptides. As shown in Fig. 2C, when 50% of the array is used to predict the other 50%, the correlation coefficient on average is well over 0.9.

### The Advantage of Analyzing Many Samples Simultaneously

The results of Fig. 3 demonstrate that simultaneous neural network analysis of all samples from all cohorts provides a somewhat more accurate overall description of binding than does sample by sample analysis. Conceptually, this suggests that there is enough information in common between the antibody molecular recognition profiles of the various samples that using the same hidden layers to describe all of them, followed by an output layer with a distinct column describing each sample, is sufficient to both describe the general and specific binding interactions. An added practical benefit to this approach is a significant reduction in computation time, as described above.

### Using the Sequence/Binding Relationship to Eliminate Noise

In Fig. 4, both the number of distinguishing peptides between cohorts and the classification accuracy improved when the measured values for each array sequence were replaced by the corresponding predicted values. Effectively, the neural network focuses information from the entire peptide dataset on each of the predicted values. This has an information aggregating effect that is extremely potent. In Fig. 5, random noise (sequence independent variation) is purposely added to the array. Since the noise is added to the log_10_ of the binding value, a sigma of 0.5 corresponds to a several-fold increase in the noise distribution width, as can be seen in Fig. 5A, and a sigma of 1 broadens the distribution of linear values by roughly an order of magnitude. As a result, multi-class classification of the original data with noise added performs poorly (Fig. 5B, dashed lines). However, because the neural network predictions effectively aggregate the combined information from nearly 123,000 sequence/binding values in the generation of the sequence/binding relationship, random noise is dramatically reduced and a sigma of 0.5 has very little effect on classification and even a sigma of 1 provides reasonable results considering that this is a 6-cohort multi-class classification problem (Fig. 5B, solid lines). This concept is taken further in Fig. 6, where arrays that for technical reasons were rejected because of excessive noise or physical artifacts affecting part of the array are included in the simultaneous analysis of all samples and their excess noise and defects are effectively repaired by comparison to other samples in the system. This is done without the network that creates the sequence-binding relationship having any information about which cohort is which in the analysis. The implication for array based diagnostic applications is that replacing a purely statistical approach like immunosignaturing with a structure-based approach provides a means of eliminating noise that is unrelated to the binding properties of the sequences (obviously, the real patient to patient variance is not removed as these differences are based on proper binding of antibodies to specific sequences).

### Using the Neural Network Model Itself for Disease Discrimination

As shown in both Fig. 4 and 6, predicted binding values for a set of peptide sequences that approximately cover the same model sequence space as the array sequences can be used to discriminate between cohorts of samples just as well as predicted values of the original array sequences. In fact, it is the sequence/binding relationship that contains the discriminating information, and it is not necessary to use predicted binding to real sequences at all. In the neural network used here for simultaneous analysis of all samples, the output layer consists of one column corresponding to each sample. The length of the column is the same as the width of the last hidden layer (350 values in this case). The 350 values associated with each sample in this output layer, combined with a single bias value added at the end, contains all of the distinguishing information for that sample and can effectively be used to replace the ∼123,000 sequence/binding values measured with only a few hundred values. Fig. 7 shows an unsupervised clustering using the algorithm UMAP(1, 3) in which the 351 values of the final weight matrix for each sample plus the bias value were used to perform a dimension reduction to 2 components. The component values for each sample are plotted and the different cohorts are color coded. The plot makes biological sense; the sera from individuals infected by viruses are clustered together but well separated into subgroups while samples from Chagas disease and uninfected individuals are distantly separated from those collected from individuals suffering viral infections. As was seen in Fig. 4, sera from WNV and HBV infected individuals are the hardest to distinguish, but the rest are almost completely distinguishable in this unsupervised analysis. Interestingly, there is one small cluster consisting of different kinds of samples completely separated from the others (upper left, Fig. 7). UMAP is a nonlinear clustering algorithm which looks for the most similar features in samples to determine clustering. Apparently, this cluster of individuals had some other unknown immunological stimulus in common that distinguished them from all others. The ability to detect such clusters could prove useful in public health bio-surveillance applications. Fig. 7 demonstrates that the cohort distinguishing information is contained in the 351 values of the final weight matrix and bias; once the sequence-binding relationship is created, there is actually no need to use predicted binding values of sequences at all in order to distinguish the different cohorts effectively.

**Fig. 7.**
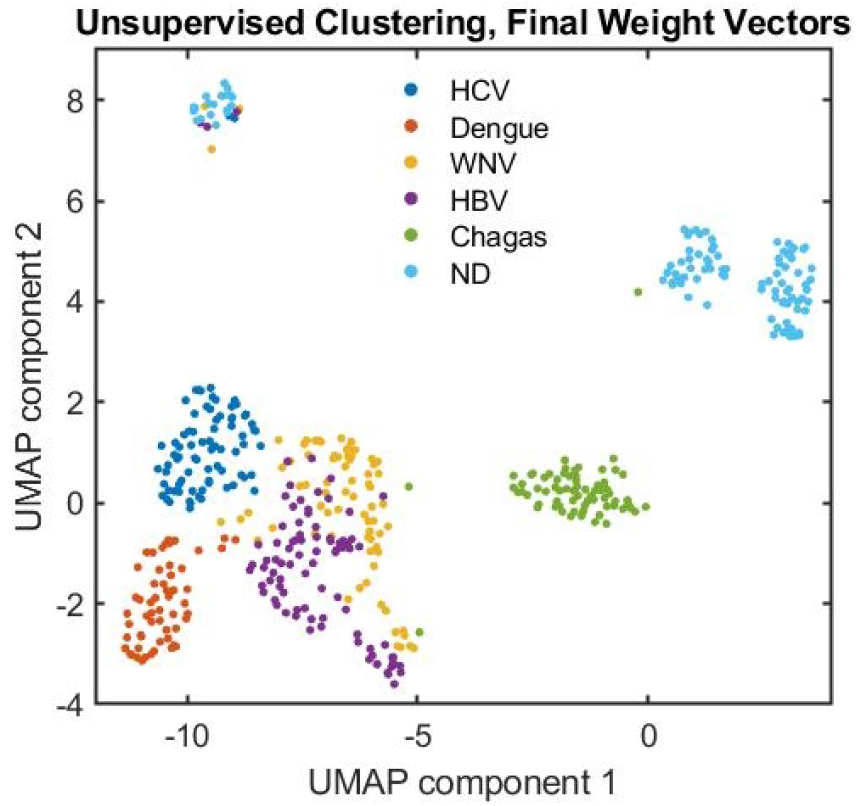
Unsupervised clustering of the neural network output layer weights. A Matlab implementation (1) of UMAP (Uniform Manifold Approximation and Projection(3)) was used to reduce 351 values from the final weight matrix of the neural network and the bias for each sample to 2 component values which are plotted. Cohorts are color coded.

## Materials and Methods

### Peptide arrays

The peptide arrays used were produced locally at ASU via photolithographically directed synthesis on silicon wafers using methods and instrumentation common in the electronics fabrication industry and as described previously(7). The synthesized wafers were cut into microscope slide sized pieces, each slide containing a total of 24 peptide arrays. Each array contained 122,926 unique peptide sequences that were 7-12 amino acids long (average of 10). A 3 amino acid linker consisting of GSG was attached to each peptide and connected the C-terminus to the array surface via amino silane. The peptides were synthesized using 16 of the 20 natural amino acids (A,D,E,F,G,H,K,L,N,P,Q,R,S,V,W,Y) in order to simplify the synthesis process (C and M were excluded due to complications with deprotection and disulfide bond formation and I and T were excluded due to the similarity with V and S and to decrease the overall synthetic complexity and the number of photolithographic steps required(45). The arrays were created in 64 photolithographic steps (4 rounds through addition of the 16 amino acids) and sequences were chosen from the set to cover all possible sequences as evenly as the synthesis would allow. A detailed description of the amino acid composition of the arrays and peptide length distribution was published previously(21) (referred to as CIMw189-S9 in that publication).

### Serum samples

Deidentified serum samples were collected from three different sources: 1) Blood donors’ samples from Creative Testing Solutions (CTS), Tempe, AZ, 2) LGC SeraCare, Milford, MA, and 3) Arizona State University (ASU) (Table 1). The dengue serotype 4 serum samples were collected from 2 of the above sources: 30 samples were provided by CTS and 35 samples were purchased by Lawrence Livermore National Labs (LLNL) from SeraCare before they were donated to the Center for Innovations in Medicine (CIM) in the Biodesign Institute at ASU. Uninfected/control samples consisted of 200 CTS samples and 18 samples from healthy volunteers at ASU. All deidentified infectious case samples came from CTS. All samples provided by CTS were residual samples collected from blood donors who were asymptomatic at the time of blood donation and were identified as test-reactive for infectious disease markers during blood screening at CTS. At the time of donation, blood donors agreed to the use of their samples in research. Serum samples were frozen shortly after collection and not thawed before being received as aliquots. ASU samples were collected under IRB protocol STUDY00002876: DHS Immunosignaturing - A Platform for Detecting and Identifying Multiple Infectious Diseases – July 2015). Serum samples were frozen at the time of collection and not thawed before being received as aliquots. Further sample description and in-house validation of disease state is described in the supplementary materials.

### Sample processing and serum IgG binding measurements

Serum from the 6 sample cohorts (5 disease cohorts and uninfected) were diluted (1:1) in glycerol and stored at -20°C. Before incubation, each serum sample was prepared as 1:625 dilution in 625 µL incubation buffer (phosphate buffered saline with 0.05 Tween 20, pH 7.2). The slides, each containing 24 separate peptide arrays were loaded into an ArrayIt microarray cassette (ArrayIt, San Mateo, CA). Then, 20 µL of the diluted serum (1:625) was added on a Whatman 903T Protein Saver Card. From the center (12 mm circle) of the protein card, a 6 mm circle was punched, and put on the top of each well in the cassette, and covered with an adhesive plate seal (3M, catalogue number: 55003076). Incubation of the diluted serum samples on the arrays was performed for 90 minutes at 37°C with rotation at 6 RPM in an Agilent Rotary incubator. Then, the arrays were washed 3 times in distilled water and dried under nitrogen. A goat anti-human IgG(H+L) secondary antibody conjugated with either AlexaFluor 555 (Life Technol.) or AlexaFluor 647 (Life Technol.) was prepared in 1x PBST pH 7.2 to a final concentration of 4 nM. Following incubation with primary antibodies, secondary antibodies were added to the array, sealed with a 3M cover and incubated at 37°C for 1 hour. Then the slides were washed 3 times with PBST (137 mM NaCl, 2.7 mM KCl, 10 mM Na_2_HPO_4_, and 1.8 mM KH_2_PO_4_. 0.1% Tween (w/v)), followed by distilled water, removed from the cassette, sprayed with isopropanol and centrifuged, dried under nitrogen, and scanned at 0.5um resolution in an Innopsys Innoscan 910 0.5 um laser scanner (Innopsys, Carbonne, Fr), excitation 547 nm, emission 590 nm. Each image was analyzed (GenePix Pro 6.0, Molecular Devices, San Jose, CA) and the raw fluorescence intensity data was exported as a GenePix Results (‘gpr’) file.

### Binding analysis using neural networks

The neural network used to relate peptide sequences on the array to the measured binding of total serum IgG has been described previously(21). The amino acid sequences are input as one-hot representations. An encoder layer linearly transforms each amino acid into a real-valued vector. The amino acid encodings are then concatenated to form a full sequence encoding. Finally, a feed-forward neural network is used to predict total serum IgG binding from the sequence encoding. The encoder and neural network are trained on the peptide sequence/binding value pairs by optimizing an L2 loss function (sum of squared error) between the measured and predicted binding values. The model performance is assessed by calculating the Pearson correlation coefficient between the measured and predicted binding values for a test dataset not involved in the training. Except where otherwise stated, the neural networks used in this work are trained on all samples simultaneously, where all layers of the encoder and neural network weights are shared across cohorts except for the final layer of the neural network.

The neural network was trained using the log_10_ of the median-normalized binding values from the peptide array (normalized by the binding values of all peptides in a given sample). Any zeros in the dataset were replaced by 0.01 x the median prior to taking the logarithm.

## Author Information

### Present Addresses

NWW, LK: Center for Molecular Design and Biomimetics, Biodesign Institute, Arizona State University, Tempe, AZ 85287

Z-GZ:: Caris Life Sciences, Tempe, AZ 85281

RC: Kriya Therapeutics, Redwood City, CA 94061

CD: Robust Diagnostics, Tempe, AZ 85287

PS: School of Life Sciences, Arizona State University, Tempe, AZ 85287

PW, VG: Creative Testing Solutions, 2424 W. Erie Dr., Tempe, AZ 85282

### Author Contributions

All authors were involved in writing or editing the manuscript. In addition:

R.C performed data analysis and conceived of approaches, A.T.T developed algorithms and concepts, L.K. developed concepts, P. S., C. D. and Z-G.Z were involved in the sample curation and data collection, N. W. W performed data analysis and conceived approaches

## Acknowledgements

The principal investigator on the DTRA grant that supported the collection of data used in this analysis was Professor Stephen Johnston; he provided valuable concepts and insights that underlie the background work upon which this analysis rests.

## Abbreviations

HCV: Hepatitis C Virus
HBV: Hepatitis B Virus
WNV: West Nile Virus
ND: Non Disease or No Known Infection
CV: Coefficient of Variation
UMAP: Uniform Manifold Approximation and Projection

## Supporting Information

**Figure S1. The correlation coefficient between the predicted and measured values for each of the 465 samples used in the analysis of Figure 2**.

## References

1. Meehan C, Ebrahimian J, Moore W, Meeha S. Uniform Manifold Approximation and Projection (UMAP) MATLAB Central File Exchange. 2022.

2. Eilers PHC, Goeman JJ. Enhancing scatterplots with smoothed densities. Bioinformatics. 2004;20(5):623–8.

3. McInnes L, Healy J, Saul N, Großberger L. UMAP: Uniform Manifold Approximation and Projection. Journal of Open Source Software. 2018;3(29):861.

4. Brown JR, Stafford P, Johnston SA, Dinu V. Statistical methods for analyzing immunosignatures. Bmc Bioinformatics. 2011;12.

5. Kukreja M, Johnston SA, Stafford P. Comparative study of classification algorithms for immunosignaturing data. Bmc Bioinformatics. 2012;13.

6. Legutki JB, Magee DM, Stafford P, Johnston SA. A general method for characterization of humoral immunity induced by a vaccine or infection. Vaccine. 2010;28(28):4529–37.

7. Legutki JB, Zhao ZG, Greving M, Woodbury N, Johnston SA, Stafford P. Scalable High-Density Peptide Arrays for Comprehensive Health Monitoring. Nat Commun. 2014;5:4785.

8. Navalkar KA, Johnston SA, Woodbury N, Galgiani JN, Magee DM, Chicacz Z, et al. Application of immunosignatures for diagnosis of valley Fever. Clin Vaccine Immunol. 2014;21(8):1169–77.

9. Nayak BP, Putterman C, Gerwien R, Sykes K, Tarasow TM. IMMUNOSIGNATURE TECHNOLOGY IDENTIFIES SYSTEMIC LUPUS ERYTHEMATOSUS FROM A DROP OF SERUM. Annals of the Rheumatic Diseases. 2016;75:1056-.

10. Restrepo L, Stafford P, Johnston SA. Feasibility of an early Alzheimer’s disease immunosignature diagnostic test. J Neuroimmunol. 2013;254(1-2):154–60.

11. Richer J, Johnston SA, Stafford P. Epitope identification from fixed-complexity random-sequence peptide microarrays. Mol Cell Proteomics. 2014.

12. Scheck AC, Stafford P, Hughes A, Cichacz Z, Coons SW, Johnston SA. Immunosignaturing for the Diagnosis and Characterization of Human Brain Tumors. Neuro-Oncology. 2012;14:100-.

13. Singh S, Stafford P, Schlauch KA, Tillett RR, Gollery M, Johnston SA, et al. Humoral Immunity Profiling of Subjects with Myalgic Encephalomyelitis Using a Random Peptide Microarray Differentiates Cases from Controls with High Specificity and Sensitivity. Mol Neurobiol. 2016.

14. Stafford P, Cichacz Z, Woodbury NW, Johnston SA. Immunosignature system for diagnosis of cancer. Proc Natl Acad Sci U S A. 2014;111(30):E3072–80.

15. Stafford P, Johnston SA, Kantarci OH, Zare-Shahabadi A, Warrington A, Rodriguez M. Antibody characterization using immunosignatures. Plos One. 2020;15(3):e0229080.

16. Sykes KF, Legutki JB, Stafford P. Immunosignaturing: a critical review. Trends Biotechnol. 2013;31(1):45–51.

17. Tarasow TM, Rowe MW, Haddad M, Sykes K. Immunosignature technology detects stage I lung cancer from a drop of serum. Cancer Research. 2015;75.

18. Rowe M, Melnick J, Gerwien R, Legutki JB, Pfeilsticker J, Tarasow TM, et al. An ImmunoSignature test distinguishes Trypanosoma cruzi, hepatitis B, hepatitis C and West Nile virus seropositivity among asymptomatic blood donors. PLoS Negl Trop Dis. 2017;11(9):e0005882.

19. Maeda D, Batista MT, Pereira LR, de Jesus Cintra M, Amorim JH, Mathias-Santos C, et al. Adjuvant-Mediated Epitope Specificity and Enhanced Neutralizing Activity of Antibodies Targeting Dengue Virus Envelope Protein. Front Immunol. 2017;8:1175.

20. Hughes AK, Cichacz Z, Scheck A, Coons SW, Johnston SA, Stafford P. Immunosignaturing Can Detect Products from Molecular Markers in Brain Cancer. Plos One. 2012;7(7).

21. Taguchi AT, Boyd J, Diehnelt CW, Legutki JB, Zhao ZG, Woodbury NW. Comprehensive Prediction of Molecular Recognition in a Combinatorial Chemical Space Using Machine Learning. ACS Comb Sci. 2020;22(10):500–8.

22. Hecker M, Fitzner B, Wendt M, Lorenz P, Flechtner K, Steinbeck F, et al. High-Density Peptide Microarray Analysis of IgG Autoantibody Reactivities in Serum and Cerebrospinal Fluid of Multiple Sclerosis Patients. Mol Cell Proteomics. 2016;15(4):1360–80.

23. Tokarz R, Mishra N, Tagliafierro T, Sameroff S, Caciula A, Chauhan L, et al. A multiplex serologic platform for diagnosis of tick-borne diseases. Scientific Reports. 2018;8(1):3158.

24. Xu GJ, Kula T, Xu Q, Li MZ, Vernon SD, Ndung’u T, et al. Viral immunology. Comprehensive serological profiling of human populations using a synthetic human virome. Science. 2015;348(6239):aaa0698.

25. Hecker M, Fitzner B, Wendt M, Lorenz P, Flechtner K, Steinbeck F, et al. High-density peptide microarray analysis of IgG autoantibody reactivities in serum and cerebrospinal fluid of multiple sclerosis patients. Molecular & cellular proteomics. 2016;15(4):1360–80.

26. Tokarz R, Mishra N, Tagliafierro T, Sameroff S, Caciula A, Chauhan L, et al. A multiplex serologic platform for diagnosis of tick-borne diseases. Scientific reports. 2018;8(1):1–10.

27. Xu GJ, Kula T, Xu Q, Li MZ, Vernon SD, Ndung’u T, et al. Comprehensive serological profiling of human populations using a synthetic human virome. Science. 2015;348(6239).

28. Ionov Y, Rogovskyy AS. Comparison of motif-based and whole-unique-sequence-based analyses of phage display library datasets generated by biopanning of anti-Borrelia burgdorferi immune sera. Plos One. 2020;15(1):e0226378.

29. Pashov A, Shivarov V, Hadzhieva M, Kostov V, Ferdinandov D, Heintz KM, et al. Diagnostic Profiling of the Human Public IgM Repertoire With Scalable Mimotope Libraries. Front Immunol. 2019;10:2796.

30. Haynes WA, Kamath K, Waitz R, Daugherty PS, Shon JC. Protein-Based Immunome Wide Association Studies (PIWAS) for the Discovery of Significant Disease-Associated Antigens. Front Immunol. 2021;12:625311.

31. Haynes WA, Kamath K, Bozekowski J, Baum-Jones E, Campbell M, Casanovas-Massana A, et al. High-resolution epitope mapping and characterization of SARS-CoV-2 antibodies in large cohorts of subjects with COVID-19. Communications Biology. 2021;4(1):1317.

32. Asif M, Orenstein Y. DeepSELEX: inferring DNA-binding preferences from HT-SELEX data using multi-class CNNs. Bioinformatics. 2020;36(Suppl_2):i634–i42.

33. Hare J, Morrison D, Nielsen M. Sampling SARS-CoV-2 Proteomes for Predicted CD8 T-Cell Epitopes as a Tool for Understanding Immunogenic Breadth and Rational Vaccine Design. Frontiers in Bioinformatics. 2021;1.

34. Hie B, Zhong ED, Berger B, Bryson B. Learning the language of viral evolution and escape. Science. 2021;371(6526):284–8.

35. Shrock E, Fujimura E, Kula T, Timms RT, Lee IH, Leng Y, et al. Viral epitope profiling of COVID-19 patients reveals cross-reactivity and correlates of severity. Science. 2020;370(6520).

36. Wu Z, Kan SBJ, Lewis RD, Wittmann BJ, Arnold FH. Machine learning-assisted directed protein evolution with combinatorial libraries. Proc Natl Acad Sci U S A. 2019;116(18):8852–8.

37. Yoshida M, Hinkley T, Tsuda S, Abul-Haija YM, McBurney RT, Kulikov V, et al. Using Evolutionary Algorithms and Machine Learning to Explore Sequence Space for the Discovery of Antimicrobial Peptides. Chem. 2018;4(3):533–43.

38. Asif M, Orenstein Y. DeepSELEX: inferring DNA-binding preferences from HT-SELEX data using multi-class CNNs. Bioinformatics. 2020;36(Supplement_2):i634–i42.

39. Hare J, Morrison D, Nielsen M. Sampling SARS-CoV-2 proteomes for predicted CD8 T-cell epitopes as a tool for understanding immunogenic breadth and rational vaccine design. Frontiers in Bioinformatics. 2021;1:1.

40. Shrock E, Fujimura E, Kula T, Timms RT, Lee I-H, Leng Y, et al. Viral epitope profiling of COVID-19 patients reveals cross-reactivity and correlates of severity. Science. 2020;370(6520).

41. Wu Z, Kan SJ, Lewis RD, Wittmann BJ, Arnold FH. Machine learning-assisted directed protein evolution with combinatorial libraries. Proceedings of the National Academy of Sciences. 2019;116(18):8852–8.

42. Yoshida M, Hinkley T, Tsuda S, Abul-Haija YM, McBurney RT, Kulikov V, et al. Using evolutionary algorithms and machine learning to explore sequence space for the discovery of antimicrobial peptides. Chem. 2018;4(3):533–43.

43. Jespersen MC, Peters B, Nielsen M, Marcatili P. BepiPred-2.0: improving sequence-based B-cell epitope prediction using conformational epitopes. Nucleic Acids Res. 2017;45(W1):W24–w9.

44. Greiff V, Redestig H, Lück J, Bruni N, Valai A, Hartmann S, et al. A minimal model of peptide binding predicts ensemble properties of serum antibodies. BMC Genomics. 2012;13(1):79.

45. Stafford P. Pseudorandom vs. Random Polymers - How to Improve the Efficiency of Lithography-Based Synthesis. 2019;1.

